# Structural insights for neutralization of BA.1 and BA.2 Omicron variants by a broadly neutralizing SARS-CoV-2 antibody

**DOI:** 10.1101/2022.05.13.491770

**Authors:** Sanjeev Kumar, Anamika Patel, Lilin Lai, Chennareddy Chakravarthy, Rajesh Valanparambil, Meredith E. Davis-Gardner, Venkata Viswanadh Edara, Susanne Linderman, Elluri Seetharami Reddy, Kamalvishnu Gottimukkala, Kaustuv Nayak, Prashant Bajpai, Vanshika Singh, Filipp Frank, Narayanaiah Cheedarla, Hans P. Verkerke, Andrew S. Neish, John D. Roback, Grace Mantus, Pawan Kumar Goel, Manju Rahi, Carl W. Davis, Jens Wrammert, Mehul S. Suthar, Rafi Ahmed, Eric Ortlund, Amit Sharma, Kaja Murali-Krishna, Anmol Chandele

## Abstract

The SARS-CoV-2 BA.1 and BA.2 (Omicron) variants contain more than 30 mutations within the spike protein and evade therapeutic monoclonal antibodies (mAbs). Here, we report a receptor-binding domain (RBD) targeting human antibody (002-S21F2) that effectively neutralizes live viral isolates of SARS-CoV-2 variants of concern (VOCs) including Alpha, Beta, Gamma, Delta, and Omicron (BA.1 and BA.2) with IC50 ranging from 0.02 – 0.05 μg/ml. This near germline antibody 002-S21F2 has unique genetic features that are distinct from any reported SARS-CoV-2 mAbs. Structural studies of the full-length IgG in complex with spike trimers (Omicron and WA.1) reveal that 002-S21F2 recognizes an epitope on the outer face of RBD (class-3 surface), outside the ACE2 binding motif and its unique molecular features enable it to overcome mutations found in the Omicron variants. The discovery and comprehensive structural analysis of 002-S21F2 provide valuable insight for broad and potent neutralization of SARS-CoV-2 Omicron variants BA.1 and BA.2.

## Main Text

The ongoing Coronavirus disease 2019 (COVID-19) pandemic caused by severe acute respiratory syndrome coronavirus 2 (SARS-CoV-2) has resulted in roughly 517 million cases and 6 million deaths worldwide (*1*). Intense global efforts are being pursued to develop, evaluate, and implement vaccines or other medical countermeasures, including monoclonal antibody (mAb) therapy (*2, 3*). Widespread transmission and key mutations have led to the emergence of viral variants that escape neutralization by therapeutic antibodies as well as natural and vaccine acquired immunity (*4–8*). Most therapeutic mAbs currently licensed for use against SARS-CoV-2 have shown reduced neutralizing activity against the Omicron (B.1.1.529) variant and its sublineages (*7–9*). This highlights a continuous need to identify mAbs that are effective against emerging variants.

Like other human coronaviruses, the spike protein of SARS-CoV-2 facilitates the entry of virus into host cells and comprises two subunits, S1 and S2 (*10*). The receptor-binding motif (RBM), a region of the receptor-binding domain (RBD) present in the S1 subunit, interacts with the host cell receptor angiotensin-converting enzyme 2 (ACE2) whereas the S2 subunit is involved in the fusion of the viral and host cell membranes (*10*). Based on their epitopes, two classification schemes have been proposed to divide RBD-specific mAbs into either four (class 1-4) or seven (RBD1-7) categories (*3, 11, 12*). An unprecedented number of mutations (>10) in the RBM of Omicron and its sublineages contribute to resistance to currently available therapeutic mAbs (*6, 7, 9, 13*).

We previously evaluated the humoral immune responses in 42 COVID-19-recovered individuals who had experienced mild symptoms after the ancestral Wuhan strain (WA.1) transmission in the year 2020 (*14*). We selected five individuals (**Table S1**) who had high SARS-CoV-2 RBD binding titers, high neutralization titers to live SARS-CoV-2 WA.1, and had detectable frequencies of RBD specific memory B cells (**Fig. S1A, S1B and S1C**) for the generation of SARS-CoV-2 RBD-specific mAbs. In total, we sorted 804 SARS-CoV-2 RBD fluorescent probe binding class-switched B cells, amplified 398 (~50%) paired heavy- and light-chain antibody gene sequences, and successfully cloned and expressed 208 antibodies (**Fig. 1A**). RBD-based ELISA screening resulted in the identification of 92 SARS-CoV-2 specific mAbs **(Fig. S1D**). These mAbs showed an average CDR3 length of 16.3 amino acids, which is typical of a human IgG repertoire (*15*) (**Fig. S1E**), as well as enriched usage of heavy and light chain variable region genes belonging to the IGHV3, IGKV1, and IGLV1 families (**Fig S1F**). Most of these mAbs had a low frequency of somatic hypermutations (SHM) in both their heavy and light chains suggesting they were recently recruited from a naive B cell pool (**Fig. S1G**). Of these mAbs, 48 blocked the ACE2-RBD interaction (**Fig. S2A**) and 18 (37.5%) successfully neutralized live virus with IC50 values ranging from 0.05 – 17 μg/ml (**Fig. S2B**). Antibody 002-S21F2 was the most potent amongst all the mAb that neutralized live SARS-CoV-2 WA.1 and thus was selected for comprehensive downstream characterization.

**Fig. 1.**
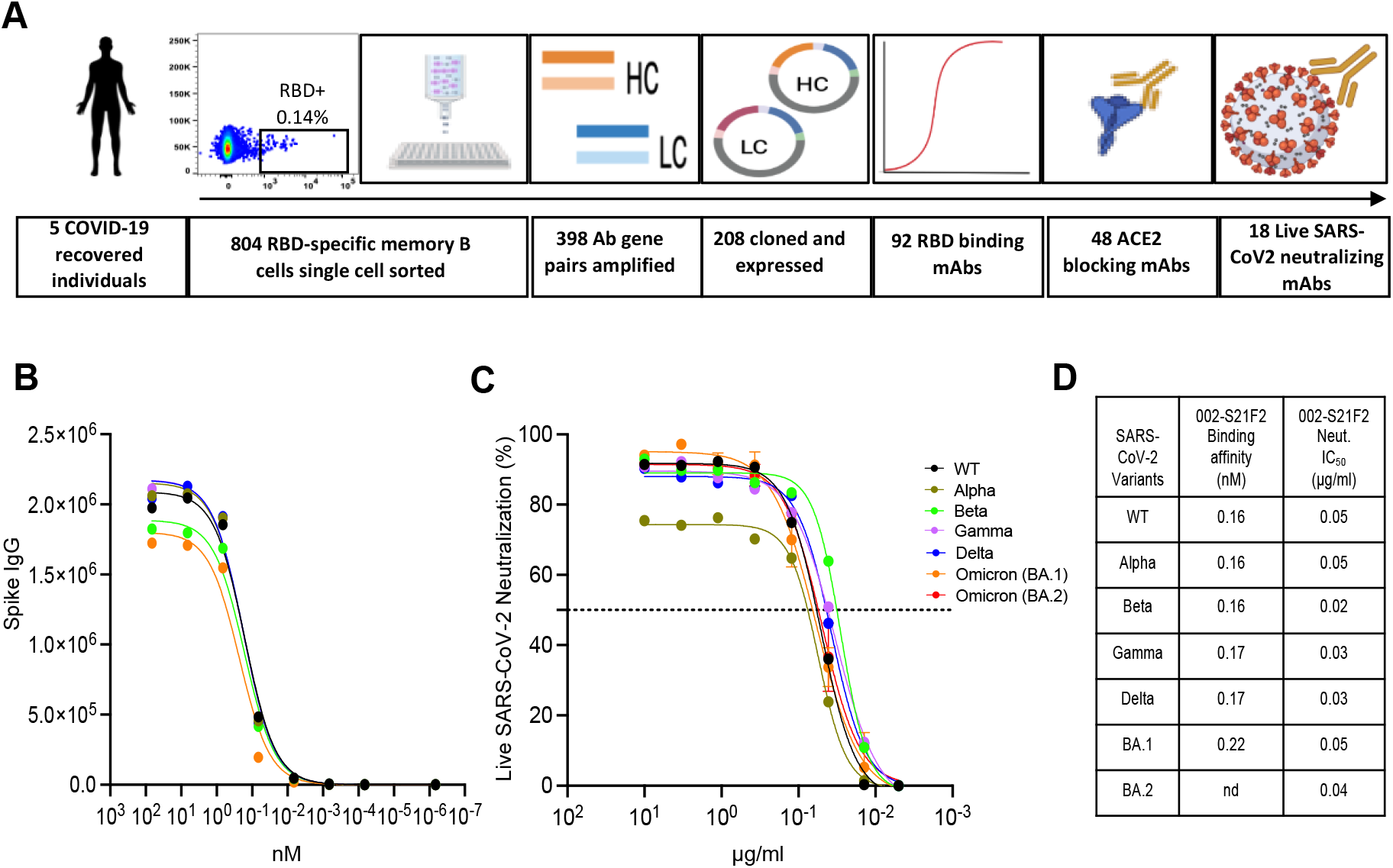
Identification of a broad and potent SARS-CoV-2 RBD specific human monoclonal antibody 002-S21F2. **(A)** The overall strategy for the isolation of RBD specific mAbs described in this study. **(B)** 002-S21F2 was tested for binding to the spike proteins of SARS-CoV-2 WA.1, Alpha, Beta, Gamma, Delta and Omicron variants of concern (VOC). **(C)** Authentic live virus neutralization curves of 002-S21F2 for WA.1, Alpha, Beta, Gamma, Delta and Omicron (BA.1 and BA.2) SARS-CoV-2 VOCs. Neutralization was determined on Vero-TMPRSS2 cells using a focus reduction assay. **(D)** 002-S21F2 mediated neutralization 50% inhibitory concentration IC50 values were obtained from live SARS-CoV-2 VOC neutralization assays. Affinity constant (KD) values calculated from the binding curves for two mAbs as measured by the MSD binding assays are plotted.

Binding analysis assessed by an electrochemiluminescence multiplex assay (Meso Scale Discovery) revealed that 002-S21F2 bound with similar affinities to all tested SARS-CoV-2 variant spike proteins including WA.1, Alpha (B.1.1.7), Beta (B.1.351), Gamma (P.1), Delta (B.1.617.2) and Omicron (B.1.1.529) (**Fig. 1B and 1D**). Furthermore, 002-S21F2 bound with picomolar affinity to the prefusion-stabilized WA.1 spike (spike-6p) by biolayer interferometry (**Fig. S3A and S3B**). Interestingly, antibody 002-S21F2 was capable of broadly neutralizing the Alpha, Beta, Gamma, Delta, BA.1 and BA.2 variants with a 50% inhibitory concentration (IC50) values of 0.05, 0.02, 0.03, 0.03, 0.05 and 0.04 μg/ml respectively (**Fig. 1C and 1D**).

To define the molecular features conferring epitope recognition and to understand the mechanism of the broad neutralization spectrum of 002-S21F2 against SARS-CoV-2 variants, we determined the cryoEM structures of 002-S21F2 full-length immunoglobulin G (IgG) in complex with WA.1 and Omicron spike-6P at 3.7 Å and 4.1 Å, respectively (**Fig. 2, S5 and S6**). The cryoEM structure showed that 002-S21F2 binds to the outer face of the RBD which is accessible in both “down” and “up” conformations and is outside the ACE2 binding motif (**Fig. 2B-C**). The interaction buried a total surface area of ~737 Å^2^ with heavy and light chains contributing ~60% and ~40% of the total interaction, respectively (**Fig. 2C**). Most of the interactions are mediated through the heavy and light-chain CDR3 regions and the epitope aligns with RBD-5/class-3 antibodies (*11, 12*). RBD residue R346 is the main contact point and is sandwiched between the heavy chain CDR3 and light chain CDR1 and CDR3 regions. Specifically, the guanidine group of R346 engages in multiple hydrogen bonds involving T102 and Y91 from the heavy and light chain CDR3, respectively (**Fig. 2C**), and has the potential for a cation-π stacking interaction involving Y32 from light chain CDR1. The T102 hydroxyl in the CDR3 heavy chain also hydrogen bonds with the RBD N448 side chain (**Fig. 2F**). The other major interaction site involves RBD residue N440, which engages in multiple interactions with W33 from CDR1 and Y52 from the heavy chain CDR2 (**Fig. 2E**). In addition, the side chain of T345 in the RBD hydrogen bonds with the main chain carbonyl of K92 in the light chain CDR3 (**Fig. 2D**). Most variants of concern (VOCs), with the exception of Omicron, do not contain any mutations within the 002-S21F2 epitope, explaining its broad neutralization ability (**Fig. 1C and Fig. 2I**). Both BA.1 and BA.2 Omicron variants contain glycine 339 to aspartic acid (G339D) and asparagine 440 to lysine (N440K) mutations within the 002-S21F2 epitope. However, the Omicron spike 002-S21F2 structure reveals identical binding compared to WA.1 and the two structures align with overall Cα-backbone RMSD of 0.975 Å and 0.875 Å in the RBD-Fab region (**Fig. S7**). All 002-S21F2 interactions observed in WA.1 remain conserved in Omicron. Furthermore, the side chain of K440 in Omicron-RBD makes an additional hydrogen bond with D57 in the heavy chain CDR2 (**Fig. 2G**), explaining why this change has minimal impact on affinity and neutralization (**Fig. 1B-D**).

**Fig. 2.**
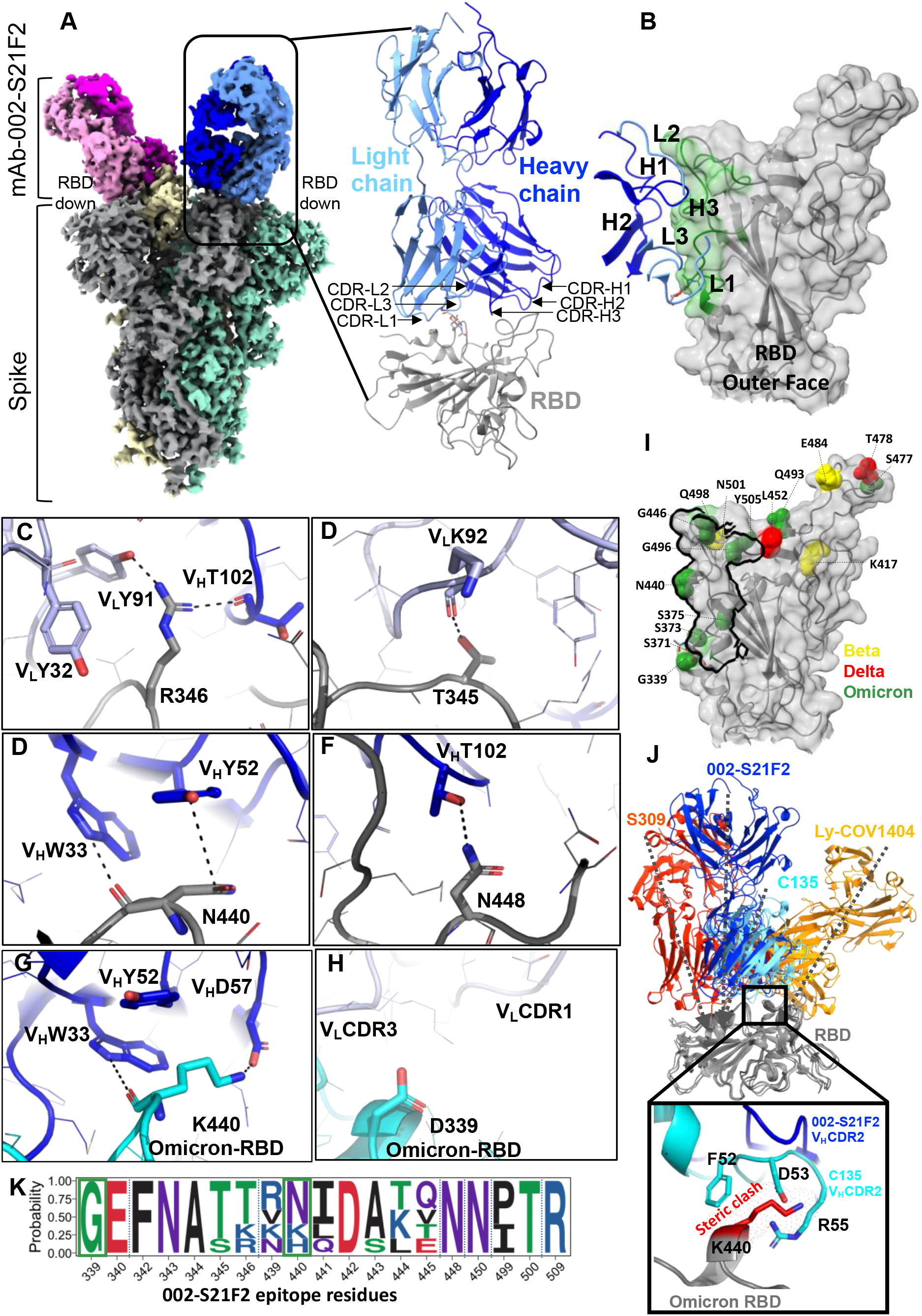
CryoEM structure of 002-S21F2 in complex with Spike trimer illustrates its binding and neutralization of VOC. **(A)** CryoEM structure of WA.1 Spike-6P trimer in complex with mAb 002-S21F2. Overall density map at contour level of 3.7 *σ* showing the antibody binding in all RBD “down” conformation. Each protomer of spike protein is shown in grey, yellow and green; the light and heavy chain of each FAB region are shown in blue/magenta and light blue/pink, respectively. The model for one of the Fab and RBD is shown in right, and the positions of all CDR regions are labelled. **(B)** Surface representation of RBD with relative positions of all CDR loops. The mapped epitope surface in RBD is highlighted in green. **(C-H)** Interaction details at 002-S21F2 and RBD binding interface, WA.1 **(C-F)** and Omicron **(G-H). (I)** locations of Beta (yellow), Delta (red) and Omicron (green) mutations on RBD relative to the 002-S21F2 epitope site that is shown as a black outline. **(J)** Structural comparison of 002-S21F2 binding mode with other class-3 mAbs, S309, C135 and Ly-COV1404; arrow representing their angle of approach on RBD. Zoomed in view showing the steric clash of Omicron K440 mutation with CDR2 residues in mAb C135. **(K)** Sequence logo representing the sequence conservation of 002-S21F2 epitope. Residue position mutated in Omicron within 002-S21F2 epitope are boxed in green. Variant mutation positions are marked above.

The antigenic residues targeted by 002-S21F2 broadly neutralizing antibody (bnAb) are highly conserved among current and previous SARS-CoV-2 VOC (**Fig. 2K and S8, S9A**). Our structural data shows that 002-S21F2 continues to maintain potent neutralization against Omicron variants BA.1 and BA.2 which harbor epitope mutations at G339D and N440K (**Fig. 1C, 1D, 2G and S7**). We suspect that this neutralization ability will persist with newly listed variants (BA.2.13, BA.2.12.1, BA.3 and BA.4/BA.5), as they are not reported to contain any additional mutations within the 002-S21F2 epitope region (Fig. **S8**). Furthermore, sequence alignment of Sarbecovirus RBDs shows 10 out of 19 conserved residues in SARS-CoV suggesting potential for cross-reactivity with other Sarbecoviruses (**Fig. 2K** and **Fig. S9B**).

Structural comparison of the 002-S21F2 epitope with other class-3 mAbs, including the two available therapeutic mAbs effective against Omicron - Ly-CoV1404 (Bebtelovimab) and S309 (Sotrovimab), show some similarities between the 002-S21F2 and C135 binding sites (**Fig. 2J**) (*16, 17*). However, C135 is unable to neutralize Omicron as a lysine mutation at RBD site N440 position would sterically clash with the C135 heavy chain CDR2 (**Fig. 2J**) (*18*). In support of this, RBD deep mutational scanning shows that an N440K mutation (present in BA.1 and BA.2) disrupts the RBD-C135 interaction (*19*).

Although 002-S21F2 recognizes an epitope outside the ACE2 binding motif, it may directly block ACE2 interaction through head-to-head inter-spike crosslinking as observed at saturating spike to IgG concentration (**Fig. S5C**). This corroborates a recent report that positively correlates high neutralization potency to inter-spike crosslinking ability within the RBD-5/class-3 antibodies (*11*). Interestingly, we also observed a higher ratio of all RBD “down” conformations (~54% particles) in antibody bound spike data compared to apo spike-6P (which only shows ~35% of all RBD “down” conformation particles). Both putative mechanisms may interfere with the ACE2 binding and contribute to neutralization.

Sequence analysis of 002-S21F2 revealed that its heavy chain (HC) variable region is comprised from VH5-51, DH5-24, and JH4 genes; the light chain (LC) gene utilizes VK1-33 and JK2 (**Fig. S4A**). Of the 5252 SARS-CoV-2 mAb sequences banked in the CoV-AbDab database (*20*), only 2 others utilized this combination of VH5-51 and VK1-33 (**Fig. S4B**). Alignment of the 002-S21F2 mAb sequence to its germline sequence revealed 4 amino acid (AA) mutations in the HC that spanned the FR1 and CDR1 regions, and 3 AA mutations present in the FR3 and CDR3 regions of the LC (**Fig. S4C**). This low frequency of somatic hypermutations (SHM), 2.7% in the HC and 1.7% in the LC, suggests that the memory B cell that expressed this mAb had not yet undergone extensive selection in the germinal center. A comparison of 002-S21F2 with SARS-CoV-2 therapeutic mAbs approved for clinical use revealed no obvious genetic similarities, suggesting that 002-S21F2 exhibits unique genetic characteristics (**Fig. 3A**).

**Fig. 3.**
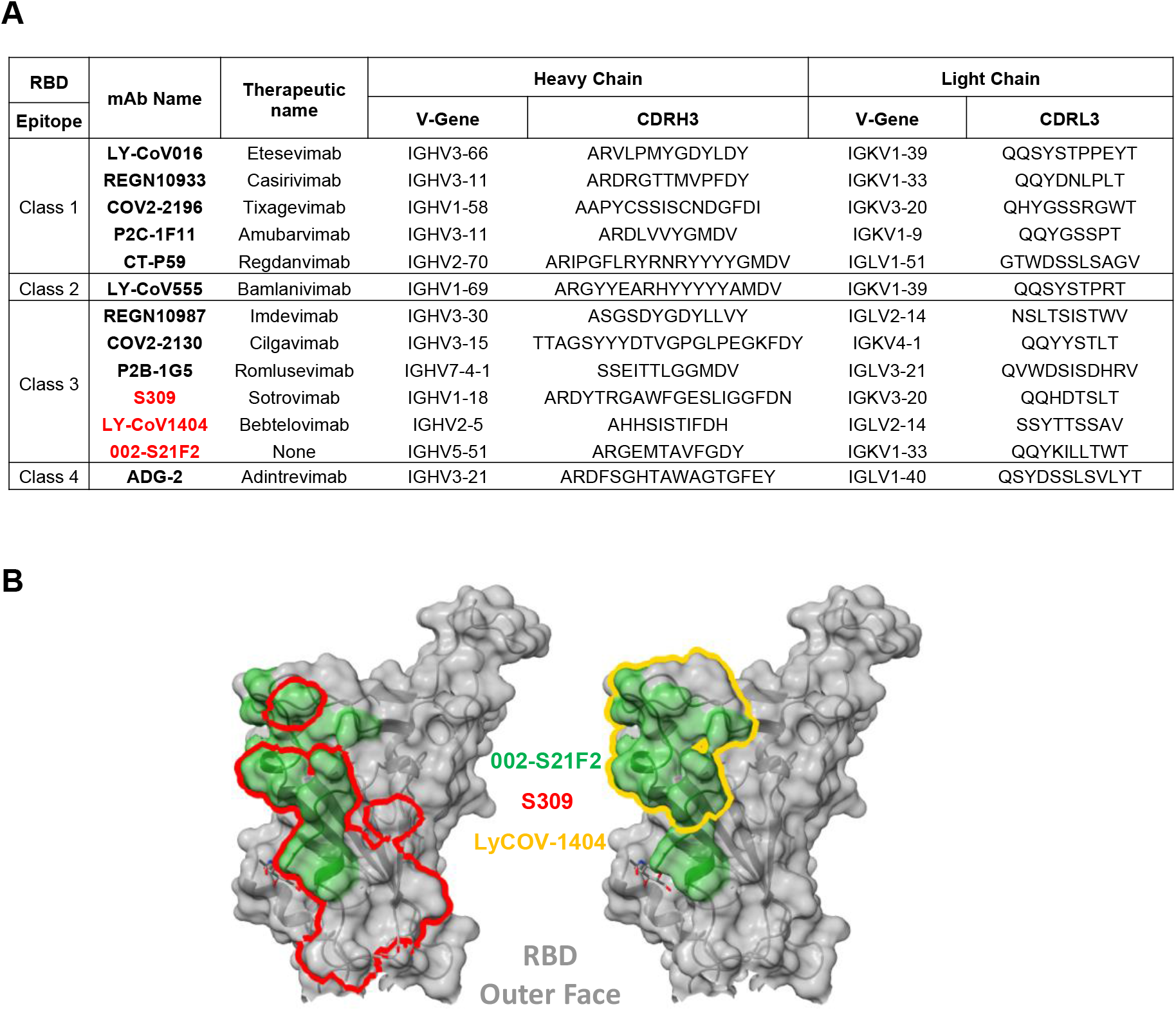
Antibody 002-S21F2 exhibits distinct genetic and epitope contact features in comparison to SARS-CoV-2 therapeutic antibodies. **(A)** Comparison of 002-S21F2 mAb genetic feature with therapeutic mAbs in clinics. Omicron neutralizing mAbs are highlighted in bold and red color **(B)** Comparison of 002-S21F2 (green) epitope site with S309 (Sotrovimab) (red outline), Ly-CoV1404 (Bebtelovimab) (yellow outline) epitopes on SARS-CoV-2 RBD.

Structural studies have reported that SARS-CoV-2 bnAbs target only a few antigenic sites on the RBD which are majorly recognized by class-3 and class-4 mAbs (*11, 12, 17, 18*). Omicron and its sublineages can evade natural and vaccine generated immunity and pose a threat to immune-compromised, vaccine-hesitant and unvaccinated adults and children. However, only 2 of the currently approved therapeutic antibodies have shown neutralization potential to Omicron – an S309 derivative (Sotrovimab) and Ly-CoV1404 (Bebtelovimab). We show that 002-S21F2 potently neutralizes both Omicron BA.1 and BA.2 and previous VOC without sacrificing potency, similar to a recently reported mAb Bebtelovimab (*16*). In contrast, Sotrovimab neutralized BA.1 with ~3-fold higher potency (than WA.1) yet poorly neutralizes BA.2 (*9*). Both of these bnAbs are class-3 antibodies that recognize overlapping epitopes on the outer face of RBD but are distinct from the 002-S21F2 epitope (**Fig. 3B)**. This suggests that the epitopes defined by the SARS-CoV-2 class-3 bnAbs target distinct antigenic residues on the outer face of the RBD, and that this surface may potentially form the basis for an effective vaccine. For example, selective steering of B cell immune responses to the RBD class-3 antigenic sites, defined by 002-S21F2 and Bebtelovimab bnAbs, may induce potent antibody responses against SARS-CoV-2 VOC. A similar successful strategy of epitope-focused vaccine candidates has been previously used to guide induction of HIV-1 bnAbs VRC01 and PGT121 that are currently in clinical trials (*21–23*). In addition, 002-S21F2 maintains potent neutralization to Omicron variants despite being isolated from a convalescent individual infected in the early months of the pandemic, when only the ancestral SARS-CoV-2 WA.1 strain was reported. Further, our structural analysis shows that the limited number of SHMs observed in this near germline bnAb 002-S21F2 are not involved in recognizing the antigenic sites (**Fig. 2**), further indicating that footprints of such bnAbs may provide a template to guide rational vaccine design.

The structural and genetic analysis of 002-S21F2 bnAb shows that it is distinct from the previously reported SARS-CoV-2 mAbs. The cryoEM structures of 002-S21F2 IgG with both WA.1 and Omicron provide a mechanistic rationale for its resilience against Omicron (BA.1 and BA.2) possibly owing to unique molecular signatures that target a non-ACE2 binding conserved RBD epitope. 002-S21F2 bnAb has tremendous potential to treat COVID-19 patients. Taken together, the discovery and structural analysis of the bnAb 002-S21F2 provides valuable insights into immune mechanisms permitting potent neutralization of highly transmissible and immune evasive Omicron VOC.

## Acknowledgements

We are thankful to Mr Satendra Singh and Mr Ajay Singh, ICGEB, New Delhi for technical support; Dr Vinay Gupta, BD Biosciences India and Aditya Rathee, ICGEB-TACF facility for single-cell sorting; Dr Vineet Menachery and Dr Pei-Yong Shi for providing the SARS-CoV-2mNG for the neutralization assays; Dr Jason McLellan for providing the SARS-CoV-2 hexapro spike expression plasmid. The cryoEM data sets on Talos Arctica were collected at Robert P. Apkarian Integrated Electron Microscopy Core (IEMC) at Emory University, Atlanta. We thank IEMC staff members for their support in data collection.

## Author contributions

Experimental work, data acquisition and analysis of data by S.K., A.P., L.L., C.R.C., R.V., M.E.D.G., V.V.E., S.L., E.S.R., K.V.G., K.N., P.B., V.S., F.F., N.C., H.V., A.S.N.., J.D.R., G.M., P.K.G., M.R., C.W.D., J.W., and E.O. Conceptualization and implementation by S.K., A.P., E.O., M.S.S., A.S., R.A., M.K.K., A.C. Manuscript writing by S.K., A.P., E.O., A.C., All authors contributed to reviewing and editing the manuscript.

## Data and materials availability

Atomic coordinates and cryoEM maps for reported structures are deposited into the Protein Data Bank (PDB) and the Electron Microscopy Data Bank (EMDB) with accession codes PDB-7U0P and EMD-26262 for WA.1 Spike-6P in complex with mAb 002-S21F2 and PDB-7UPL and EMD-26669 for Omicron Spike-6P in complex with mAb 002-S21F2. Immunoglobulin sequences are available in GenBank under accession numbers XX. Any additional data are available upon reasonable request from the corresponding authors. Source data are provided in this paper.

## Materials and Methods

### Human subjects

COVID-19 recovered individuals have been described earlier (*14*). Of these, five subjects chosen based on the frequency of receptor binding protein-positive memory B cells and the available number of banked peripheral blood mononuclear cells (PBMCs) were included in this study for human monoclonal antibodies generation.

### SARS-CoV-2 RBD-specific ELISA binding assays

The recombinant SARS-CoV-2 RBD gene was cloned, expressed, purified and ELISAs were performed as previously described (*24*). Briefly, purified RBD was coated on MaxiSorp plates (Thermo Fisher, #439454) at a concentration of 1 μg/mL in phosphate-buffered saline (PBS) at 4°C overnight. The plates were washed extensively with PBS containing 0.05% Tween-20. Three-fold serially diluted plasma or purified mAb was added to the plates and incubated at room temperature for 1 hr. After incubation, the plates were washed and the SARS-CoV-2 RBD specific IgG, IgM, IgA signals were detected by incubating with horseradish peroxidase (HRP) conjugated - anti-human IgG (Jackson ImmunoResearch Labs, #109-036-098), IgM (Jackson ImmunoResearch Labs, #109-036-129), or IgA (Jackson ImmunoResearch Labs, #109-036-011). Plates were then washed thoroughly and developed with o-phenylenediamine (OPD) substrate (Sigma, #P8787) in 0.05M phosphate-citrate buffer (Sigma, #P4809) pH 5.0, containing 0.012% hydrogen peroxide (Fisher Scientific, #18755). Absorbance was measured at 490 nm.

### Authentic live SARS-CoV-2 neutralization assay

Neutralization titers to SARS-CoV-2 were determined as previously described (*14, 24*). Briefly, 100 pfu of SARS-CoV-2 (2019-nCoV/USA_WA1/2020), Alpha, Beta, Gamma, Delta and Omicron (BA.1 and BA.2) were used on Vero TMPRSS2 cells. Heat-inactivated serum (only for WT) or purified monoclonal was serially diluted three-fold in duplicate starting at a 1:20 dilution or 10 μg/ml respectively in a 96-well round-bottom plate and incubated for 1 h at 37°C. This antibody-virus mixture was transferred into the wells of a 96-well plate that had been seeded with Vero-TMPRSS2 cells the previous day at a concentration of 2.5 × 10^4^ cells/well. After 1 hour, the antibody-virus inoculum was removed and 0.85% methylcellulose in 2% FBS containing DMEM was overlaid onto the cell monolayer. Cells were incubated at 37°C for 16-40 hours. Cells were washed three times with 1X PBS (Corning Cellgro) and fixed with 125 μl of 2% paraformaldehyde in PBS (Electron Microscopy Sciences) for 30 minutes. Following fixation, plates were washed twice with PBS and 100 μl of permeabilization buffer, was added to the fixed cells for 20 minutes. Cells were incubated with an anti-SARS-CoV spike primary antibody directly conjugated with alexaflour-647 (CR3022-AF647) for up to 4 hours at room temperature.

Plates were then washed twice with 1x PBS and imaged on an ELISPOT reader (CTL Analyzer). Foci were counted using Viridot (counted first under the “green light” set followed by background subtraction under the “red light” setting). IC50 titers were calculated by non-linear regression analysis using the 4PL sigmoidal dose curve equation on Prism 9 (Graphpad Software). Neutralization titers were calculated as 100% x [1-(average foci in duplicate wells incubated with the specimen) ÷ (average number of foci in the duplicate wells incubated at the highest dilution of the respective specimen).

### SARS-CoV-2 RBD specific memory B cell staining and single-cell sorting

Purified SARS-CoV-2 RBD protein was labelled with Alexa Fluor 488 using a microscale protein labelling kit (Life Technologies, #A30006) as per the manufacturer’s protocol. Ten million PBMCs of select COVID-19 recovered donors were stained with RBD-Alexa Fluor 488 for 1 hour at 4°C, followed by washing with PBS containing 2% FBS (FACS buffer) and incubation with efluor780 Fixable Viability (Live Dead) dye (Life Technologies, #65-0865-14) and anti-human CD3, CD19, CD20, CD27, CD38 and IgD antibodies (BD Biosciences) for 30 minutes. Cells were washed twice with FACS buffer and acquired on BD FACS ARIA Fusion (BD Biosciences). Live IgD negative B cells that were positive for the SARS-CoV-2 RBD-Alexa Fluor 488 protein were single cell sorted into a 96-well plate containing 5 μl of lysis buffer. The lysis buffer consisted of 20 U of RNase inhibitor (Promega), in 10 mM Tris pH 8.0 buffer. The plates with the sorted single cells were centrifuged gently at 2000 rpm at 4°C and stored immediately at −80°C for at least 1 hr before performing the cDNA synthesis. Data were analyzed using FlowJo software 10.

### Antibody genes amplification and cloning

The antibody genes were amplified as described earlier (*25, 26*). Briefly, cDNA was synthesized, and antibody variable gene VDJ segments were amplified by reverse transcription-polymerase chain reaction (RT-PCR) using a template-switching rapid amplification of complementary DNA (cDNA) ends (RACE) approach (Davis et al., manuscript in preparation). Gene segments were cloned into AbVec6W vectors (*25*). 4 colonies from each transformed plate were randomly picked and the insert was checked by performing colony PCR using nested PCR primers. The sequence integrity of the plasmids was verified by Sanger sequencing (Macrogen sequencing, South Korea).

### Immunogenetic analyses of antibody genes

The immunogenetic analysis of both heavy chain and light chain germline assignment, framework region annotation, determination of somatic hypermutation (SHM) levels (nucleotides) and CDR loop lengths (amino acids) was performed with the aid of IMGT/HighV-QUEST (www.imgt.org/HighV-QUEST) (*27*).

### Expression of human monoclonal antibodies

For small-scale transfection, expi293F cells were maintained in 293 expression medium and transfected at a density of 2.5 million cells per/ml in a volume of 4 ml culture per well of a 6-well cell culture plate (Corning). The transfection mix consisted of a 1:1.5 HC/LC ratio using a 1:3 ratio with 1 mg/ml PEI-Max transfection reagent (Polysciences) in 200 μl Opti-MEM. After 15-minute incubation at RT, the transfection mix was added to the cells. Supernatants were harvested 4-5 days post-transfection and clarified supernatants were tested for their SARS-CoV-2 RBD binding potential by enzyme-linked immunosorbent assay (ELISA). Supernatant with positive RBD binding signals was next purified using Protein A/G beads (Thermo Scientific), concentrated using a 30 kDa or 100 kDa cut-off concentrator (Vivaspin, Sartorius) and stored at 4°C for further use.

### SARS-CoV-2 surrogate virus neutralization test (sVNT)

The potential of human ACE2 and SARS-CoV-2 RBD interaction inhibition by RBD-specific mAbs was measured with the cPass SARS-CoV-2 surrogate virus neutralization test (sVNT) kit (Genscript, Singapore) as described previously (*28*), as per the manufacturer’s protocol. Briefly, each mAb at 20 μg/ml concentration was mixed with equal volumes of recombinant HRP-conjugated RBD and incubated for 30 min at 37°C. Next, 100 μl of this mixture was transferred to 96-well plates coated with recombinant hACE2 receptor and further incubated for 15 min at 37°C. The plate was washed four times with 1X PBST buffer followed by the addition of tetramethylbenzidine (TMB) substrate). The plate was incubated for 15 min at room temperature, and the reaction was stopped by adding the stop solution. Absorbance was measured at 450 nm and the percentage of inhibition of each sample was calculated using the following formula: % inhibition = (1- (OD450 sample/ OD450 of negative control)) x 100. Controls were included in duplicate; samples were analyzed in the singular. Inhibition >30% was regarded as a positive neutralization.

### Electrochemiluminescence antibody binding assay

Binding analysis of SARS-CoV-2 mAb to spike protein was performed using an electrochemiluminescence assay as previously described (*29*). V-PLEX COVID-19 Panel 24 (Meso Scale Discovery) was used to measure the IgG1 mAb binding to SARS-CoV-2 spike antigens following the manufacturer’s recommendations. Briefly, antigen coated plates were blocked with 150 μl/well of 5% BSA in PBS for 30 minutes. Plates were washed three times with 150 μl/well of PBS with 0.05% Tween between each incubation step. mAbs were serially diluted for concentrations ranging from 10 μg/ml to 0.1 pg/ml and 50 μl/well were added to the plate and incubated for two hours at room temperature with shaking at 700rpm. mAb antibody binding was then detected with 50 μl/well of MSD SULFO-TAG anti-human IgG antibody (diluted 1:200) incubated for one hour at room temperature with shaking at 700rpm. 150 μl/well of MSD Gold Read Buffer B was then added to each plate immediately before reading on an MSD QuickPlex plate reader.

### Octet BLI analysis

Octet biolayer interferometry (BLI) was performed using an Octet Red96 instrument (ForteBio, Inc.). A 5 μg/ml concentration of 002-S21F2 was captured on a protein A sensor and its binding kinetics were tested with serial 2-fold diluted RBD (600 nM to 37.5 nM) and spike hexapro protein (100 nM to 6.25 nM). The baseline was obtained by measurements taken for 60 s in BLI buffer (1x PBS and 0.05% Tween-20), and then, the sensors were subjected to association phase immersion for 300 s in wells containing serial dilutions of RBD or trimeric spike hexapro protein. Then, the sensors were immersed in BLI buffer for as long as 600 s to measure the dissociation phase. The mean K_on_, K_off_ and apparent KD values of the mAbs binding affinities for RBD and spike hexapro were calculated from all the binding curves based on their global fit to a 1:1 Langmuir binding model using Octet software version 12.0.

### Spike protein expression and purification

SARS-CoV-2 Spike-6P trimer protein carrying WA.1 and Omicron strain mutations were produced by transfecting FreeStyle 293-F cells using WA.1-spike-6P and Omicron-spike-6P DNA plasmids, respectively. There are two mismatched positions in our Omicron plasmid −1) position 213-216 in NTD is EPER instead of sequence REPE, 2) position 493 is a lysine residue instead of an arginine. Transfections were performed as per the manufacturer’s protocol (Thermo Fisher). Briefly, FreeStyle 293-F cells were seeded at a density of 2×10^6^ cells/ml in Expi293 expression medium and incubated at 37°C and 127 rpm with 8% CO2 overnight. The next day, 2.5×106 cells/ml were transfected using ExpiFectamineTM 293 transfection reagent (ThermoFisher, cat. no. A14524). The cells continued to grow for 4-5 days at 37°C, 127 rpm, 8% CO2 incubator. The cells were removed by centrifugation at 4,000g for 20 minutes at room temperature and spike protein-containing supernatant was collected. The supernatant was filtered and loaded onto pre-washed His-Pur Ni-NTA resin for affinity purification. The Ni-NTA resin was incubated with a spike-trimer containing supernatant for 2 hours on a shaker at room temperature. Resin washed with wash Buffer containing 25mM Imidazole, 6.7mM NaH2PO4.H2O and 300mM NaCl in PBS followed by spike protein elution in elution buffer containing 235mM Imidazole, 6.7mM NaH2PO4.H2O and 300mM NaCl in PBS. Eluted protein dialyzed against PBS and concentrated. The concentrated protein ran onto a Superose-6 Increase 10/300 column and protein eluted as trimeric spike collected. The quality of the protein was evaluated by SDS-PAGE and by Negative Stain-EM.

### Negative Stain – Electron Microscopy (NS-EM)

Spike protein was diluted to 0.05mg/ml in PBS before grid preparation. A 3μL drop of diluted protein was applied to previously glow-discharged, carbon-coated grids for ~60 sec, blotted and washed twice with water, stained with 0.75% uranyl formate, blotted and air-dried. Between 30-and 50 images were collected on a Talos L120C microscope (Thermo Fisher) at 73,000 magnification and 1.97 Å pixel size. Relion-3.1 (*30*) or Cryosparc v3.3.2 (*31*) was used for particle picking, 2D classification.

### Sample preparation for cryoEM

SARS-CoV-2 spike-6P trimer incubated with the mAb (full-length IgG) at 0.7 mg/ml concentration. The complex was prepared at a 0.4 sub-molar ratio of mAb to prevent inter-spike crosslinking, mediated by bi-valent binding of Fab in IgG. The complex was incubated at room temperature for ~5 min before vitrification. 3μL of the complex was applied onto a freshly glow-discharged (PLECO easiGLOW) 400 mesh, 1.2/1.3 C-Flat grid (Electron Microscopy Sciences). After 20 s of incubation, grids were blotted for 3 s at 0 blot force and vitrified using a Vitrobot IV (Thermo Fisher Scientific) under 22°C with 100% humidity.

### CryoEM data acquisition

Single-particle cryoEM data for mAb 002-S21F2 in complex with WA.1 and Omicron spike-6p protein were collected on a 200 kV Talose Arctica transmission electron microscope (ThermoFisher Scientific) equipped with Gatan K3 direct electron detector behind 20 eV slit width energy filter. Multi-frame movies were collected at a pixel size of 1.1 Å per pixel with a total dose of 51 e/Å^2^ at defocus range of −1.0 to −2.4 μm.

### CryoEM data analysis and model building

CryoEM movies were motion-corrected either in Motioncorr2 in Relion3.0 (*30*) or using Patch motion correction implemented in Cryosparc v3.3.1 (*31*). When Motion correction was performed outside of Cryosparc, motion-corrected micrographs were imported in Cryosparc v3.3.1 and corrected for contrast transfer function using Cryosparc’s implementation of Patch CTF estimation. Micrographs with poor CTF fits were discarded using CTF fit resolution cutoff to ~6.0 Å. Particles were picked using a Blob picker, extracted and subjected to an iterative round of 2D classification. Particles belonging to the best 2D classes with secondary structure features were selected for heterogeneous 3D refinement to separate IgG bound Spike particles from non-IgG bound Spike particles. Particles belonging to the best IgG bound 3D class were refined in non-uniform 3D refinement with per particle CTF and higher-order aberration correction turned on. To further improve the resolution of the RBD-IgG binding interface a soft mask was created covering one RBD and interacting Fab region of IgG and refined locally in Cryosparc using Local Refinement on signal subtracted particles. All maps were density modified in Phenix (*32*) using Resolve CryoEM. The combined Focused Map tool in Phenix was used to integrate high resolution locally refined maps into an overall map. Additional data processing details are summarized in Figure **S5**-**S6**.

The initial spike models for WA.1 (PDB:7lrt) or Omicron (PDB:7tf8) as well as individual heavy and light chains of the Fab region of an IgG (generated with Alphafold (*33*)) were docked into cryoEM density maps using UCSF ChimeraX (*34*). The full Spike-mAb model was refined using rigid body refinement in Phenix, followed by refinement in Isolde (*35*). The final model was refined further in Phenix using real-space refinement. Glycans with visible density were modelled in Coot (*36*). Model validation was performed using Molprobity (*37*). PDBePISA (*38*) was used to identify mAb-RBD interface residue, to calculate buried surface area and to identify polar interaction. Figures were prepared in ChimeraX(*34*) and PyMOL (*39*).

## Supplementary Figures and Tables

**Fig. S1:**
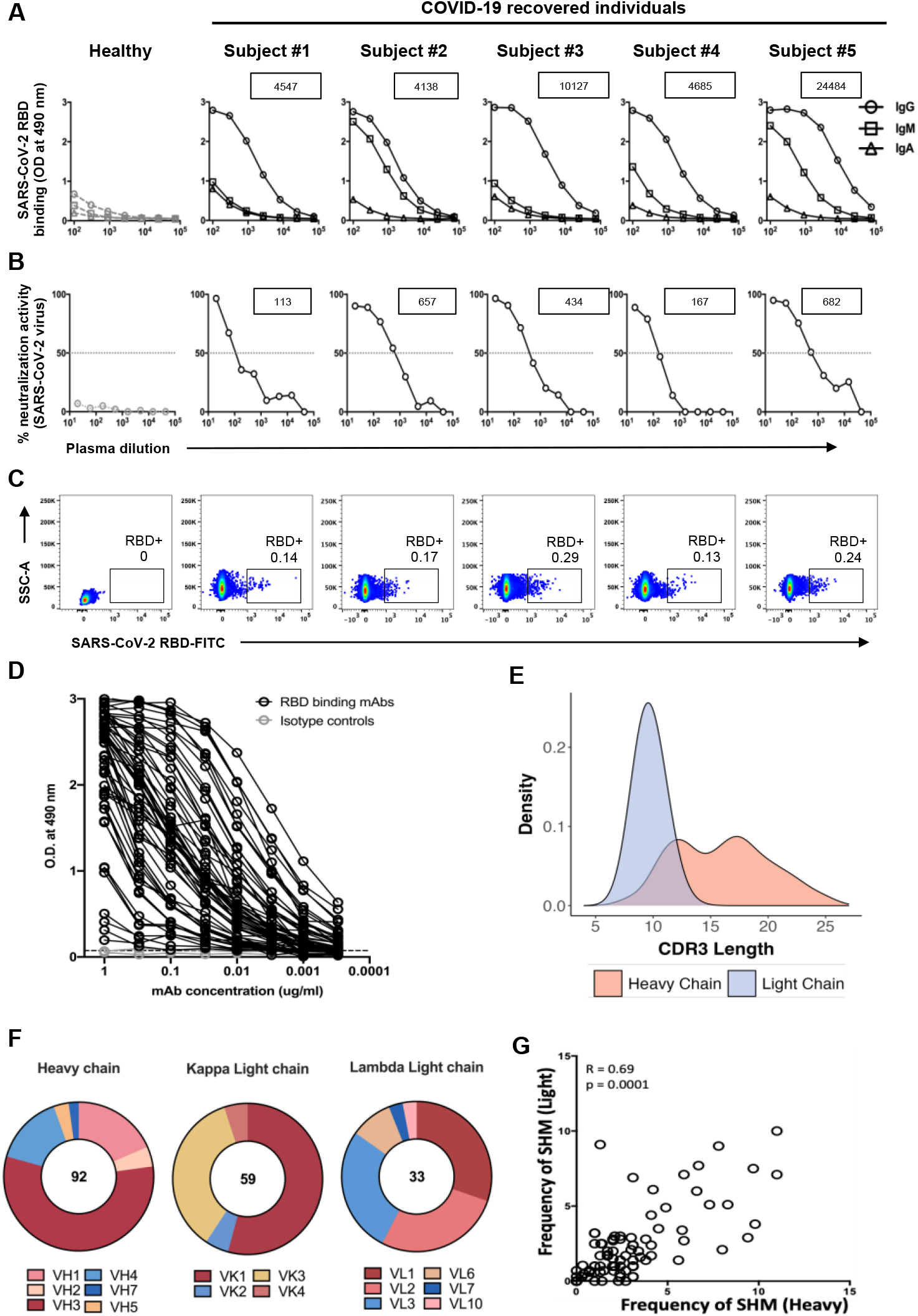
Isolation and characterization of 92 SARS-CoV-2 RBD specific mAbs. **(A)** ELISA curves showing plasma IgG, IgM and IgA reactivity against SARS-CoV-2 RBD protein. The five subjects are COVID-19 recovered individuals recruited into a clinical study during the first wave of the pandemic in India. ELISA results with the plasma of a pre-pandemic healthy individual are shown for comparison. **(B)** Neutralization assay curves showing activity against SARS-CoV-2 WA1/2020 strain and Delta (B.1.617.2) strain. The plasma dilution resulting in a 50% reduction in neutralization (FRNT-mNG_50_) is indicated by arrows. **(C)** Flow cytometry plots show SARS-CoV-2 RBD-specific memory B cells in the peripheral blood lymphocytes of select SARS-CoV-2 recovered individuals. The plots shown were gated on CD19 positive and CD3 negative populations. Staining was performed using FITC conjugated RBD protein. **(D)** ELISA curves showing SARS-CoV-2 RBD binding by each of the purified mAbs (n=92). **(E)** The CDR3 length of each sequence was calculated and their distribution was plotted as a histogram. **(F)** Donut plots showing the heavy chain, lambda light chain and kappa light chain gene distribution of the SARS-CoV-2 RBD-specific mAbs (n=92). **(G)** Correlation plot of heavy chain and light chain SHM (%). Pearson correlation coefficient (R) and p-value have been shown.

**Fig. S2:**
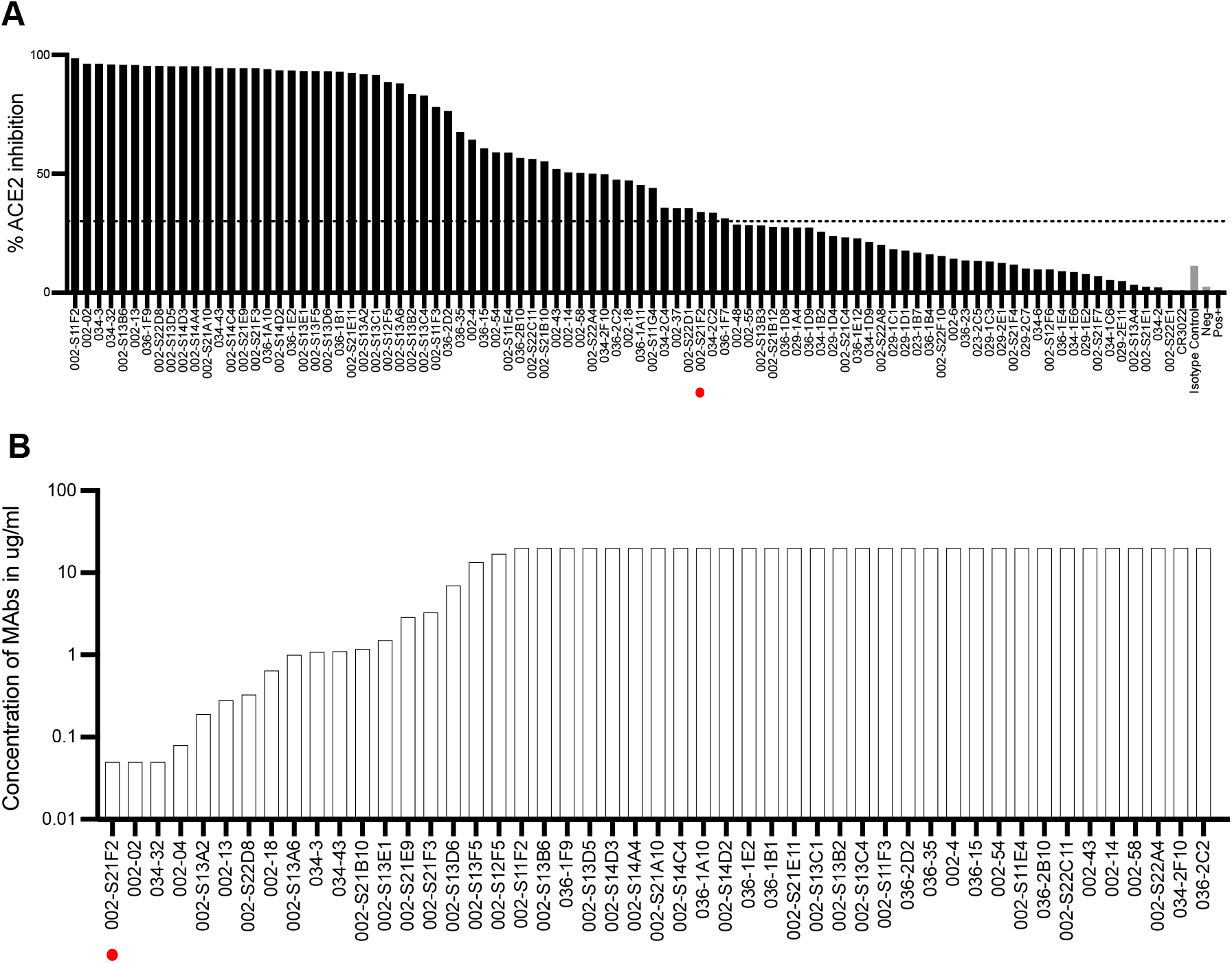
ACE2 inhibition and neutralization potential analysis of 92 SARS-CoV-2 RBD specific mAbs. **(A)** ACE2 inhibition analysis of SARS-CoV-2 RBD specific human monoclonal antibodies (n=92) was performed using the cPass™ SARS-CoV-2 surrogate neutralization antibody detection kit (Genscript, USA). All mAbs were tested at 20 μg/ml. Percent (%) inhibition values are plotted as a bar diagram (Black colour). Here, CR3022, a SARS-CoV and SARS-CoV-2 specific mAb was used as a positive control, A5GK (an inhouse CHIKV specific mAb) and D2-DV (an inhouse DENV mAb) were used as negative isotype control mAbs. The dotted line of the 30% cut-off value shows the set threshold over which ACE2 inhibition is positive as per manufacturers’ instructions. **(B)** Neutralization titers (FRNT-mNG_50_) against SARS-CoV-2 live virus (stain (WA.1/2020) of the few RBD binding mAbs. Here, mAb 002-S21F2 is marked with a red dot.

**Fig. S3:**
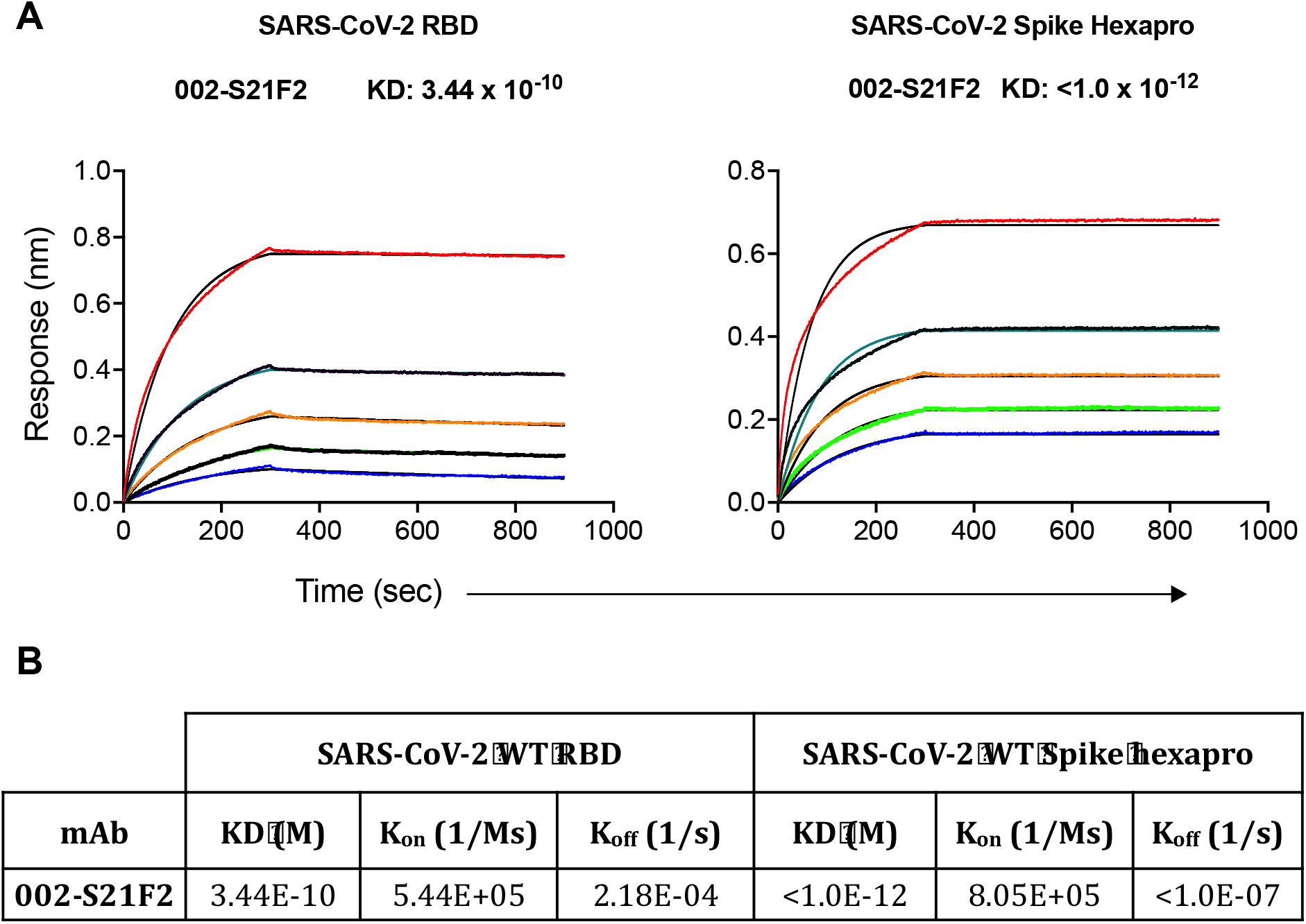
Antibody 002-S21F2 showed high affinity to SARS-CoV-2 RBD and spike proteins in biolayer interferometry assays. **(A)** Octet BLI sensorgrams showing the SARS-CoV-2 RBD and Spike-6p binding affinities of the mAb 002-S21F2. In these assays, 002-S21F2 mAb (5 ug/ml concentration) was captured on protein A sensors and its binding kinetics were tested with serial 2-fold diluted RBD (600 nM to 37.5 nM) and Spike hexapro protein (100 nM to 6.25 nM). Association was measured for 300 seconds followed by dissociation measurement for 600 seconds. **(B)** Describing the KD (M), K_on_ (1/Ms) and K_off_ (1/s) values of the four potent mAbs with RBD and Spike hexapro proteins.

**Fig. S4.**
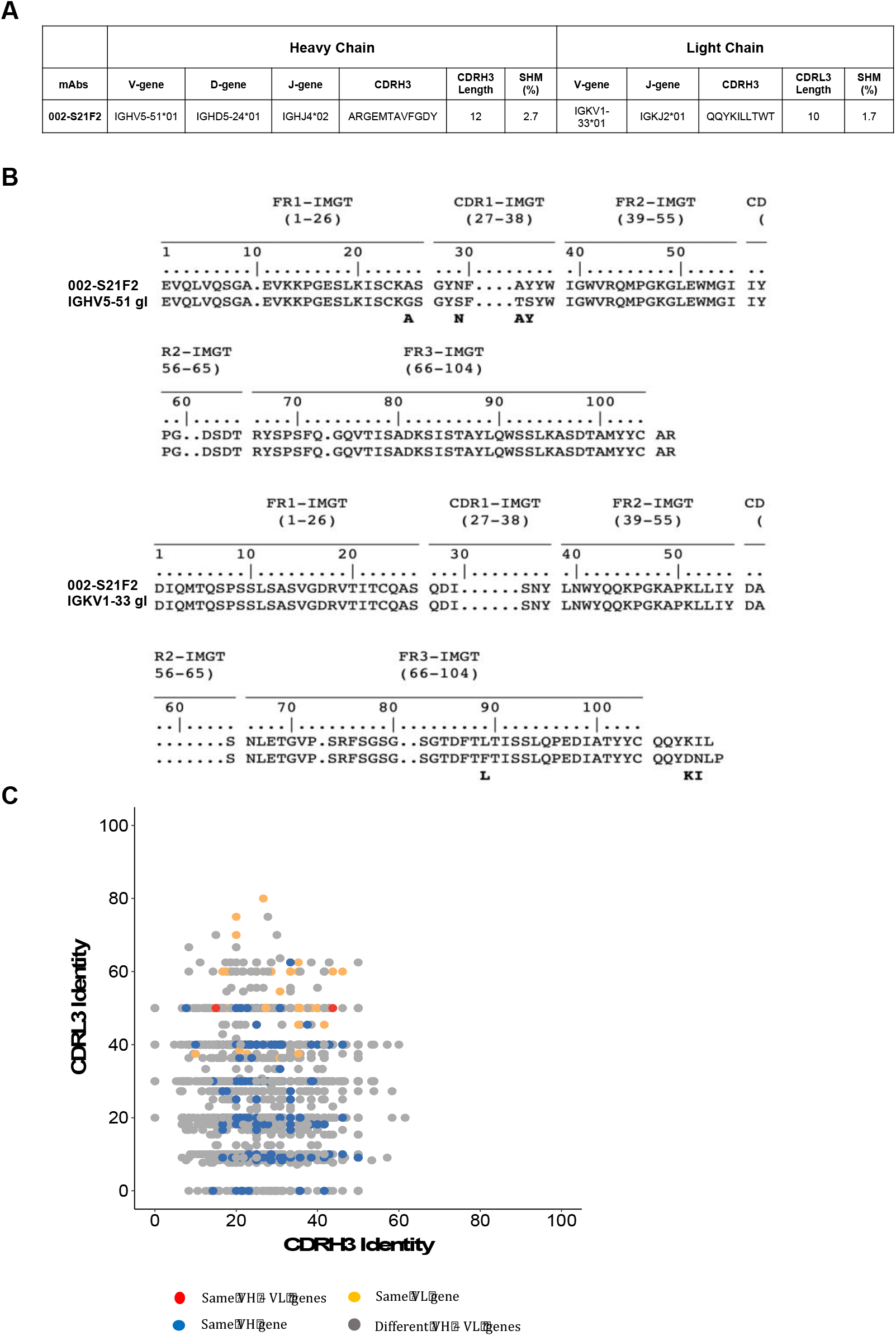
SARS-CoV-2 RBD specific bnAb 002-S21F2 exhibits a rare immunogenetic make-up. **(A)** Table showing the antibody heavy chain (HC) and light chain (LC) genetic information of the 002-S21F2 bnAb. **(B)** IMGT V-quest alignment analysis result of 002-S21F2 HC and LC gene with their respective germline sequence. Here, somatic mutations are marked as bold amino acids in black color. **(C)** CDRH3 amino acid sequence identity to 002-S21F2 is plotted against CDRL3 amino acid identity to 002-S21F2 for paired HC and LC sequences of SARS-CoV-2 mAbs banked in the CoV-AbDab database. SARS-CoV-2 mAbs with the same HC and LC germline gene as 002-S21F2 (IGHV5-51 and IGKV1-33) are shown in red color. MAbs using the same HC gene but different LC gene as 002-S21F2 are shown in blue color. MAbs using different HC genes but the same LC gene as 002-S21F2 are shown in orange color. MAbs using a different HC and LC combination as 002-S21F2 are shown in grey color.

**Fig. S5.**
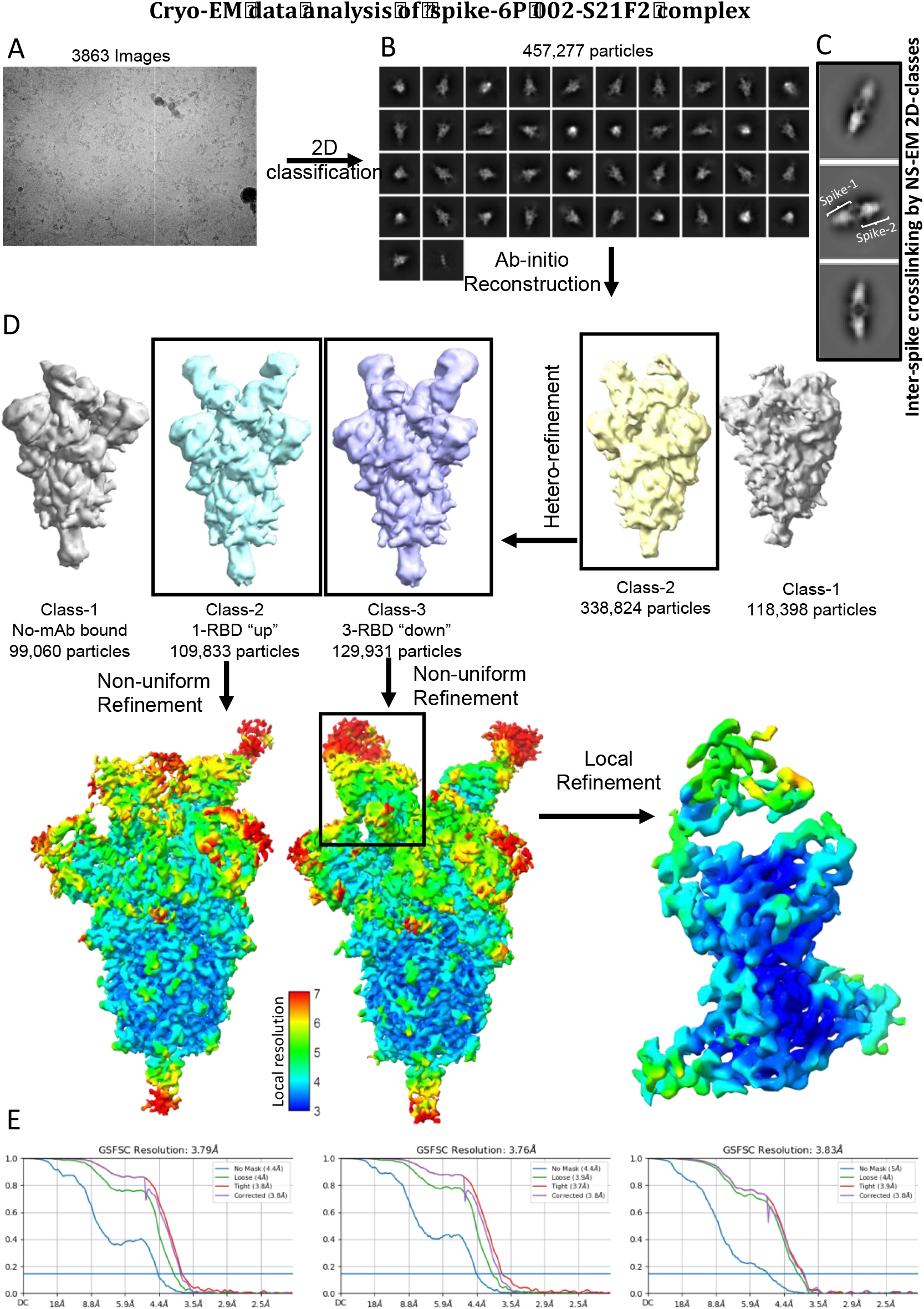
CryoEM data analysis and validation for WA.1 Spike-6P 002-S21F2 complex. **(A)** Representative electron micrograph. **(B)** Representative 2D-class averages. **(C)** The 2D-Classes derived from Negative Stain EM (NS-EM) data of WA.1 Spike-6P and 002-S21F2 complex at saturating IgG concentration show head-to-head spike dimer. **(D)** Classification scheme and refinement that yielded final cryoEM map reconstruction. Boxed classes were selected for further processing and refinement. The boxed region contains one RBD complexed with one Fab in a refined map masked for local refinement**. (E)** Gold standard Fourier shell correlation curve of final overall (two left panels) and locally refined (right panel) maps and resolution estimation based on 0.143 Fourier shell correlation criteria as indicated by a blue line.

**Fig. S6.**
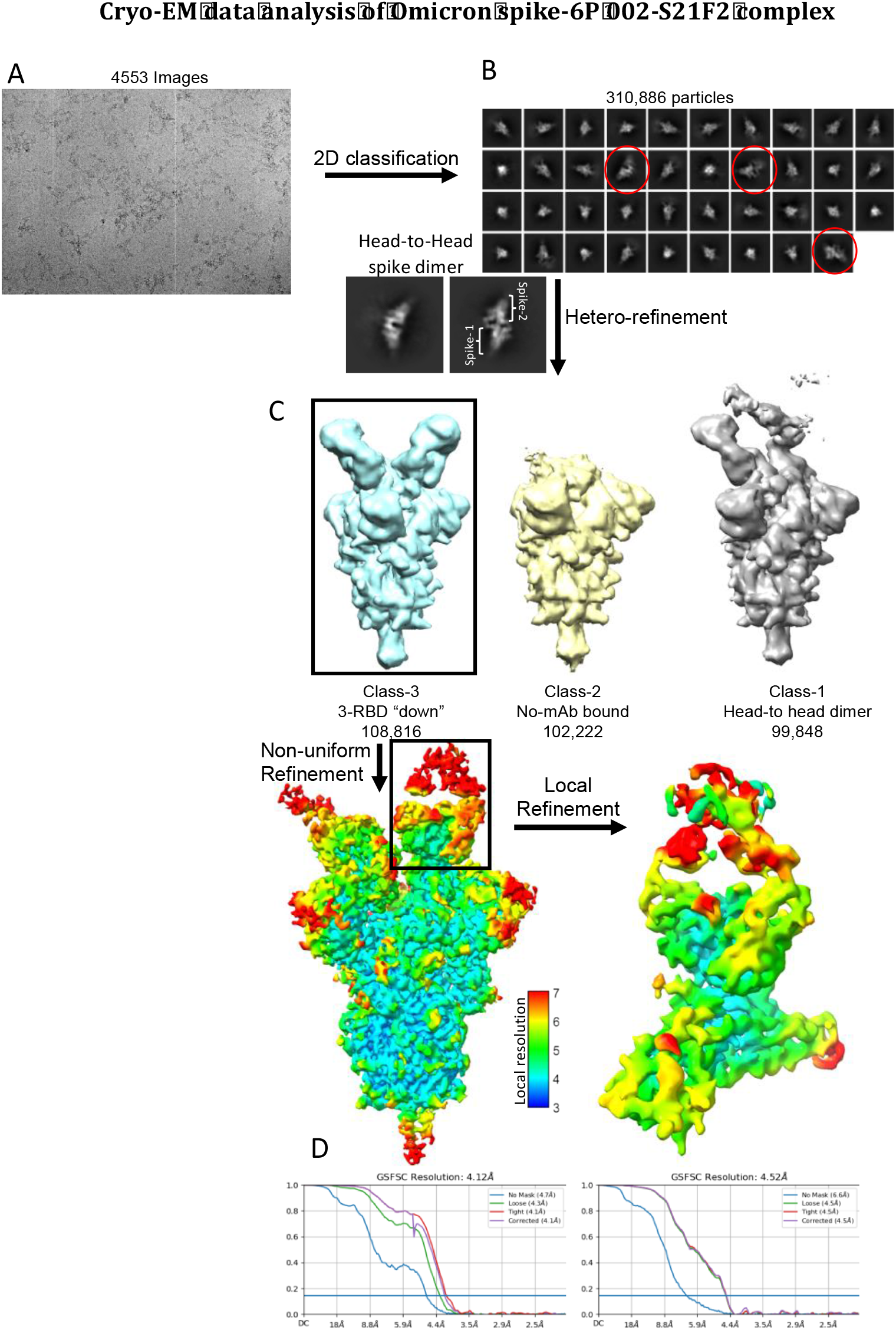
Cryo-EM data analysis and validation for Omicron Spike-6P and 002-S21F2 complex. **(A)** Representative electron micrograph. **(B)** Representative 2D-class averages. 2D classes showing head-to-head spike dimer are circled in red and shown below. **(C)** Classification scheme and refinement that yielded final cryo-EM map reconstruction. Boxed classes were selected for further processing and refinement. Boxed region containing one RBD complexed with one Fab in refined map masked for local refinement. **(D)** Gold standard Fourier shell correlation (FSC) curve of final overall (left) and locally refined (right) maps and resolution estimation based on 0.143 Fourier shell correlation criteria as indicated by a blue line.

**Fig. S7.**
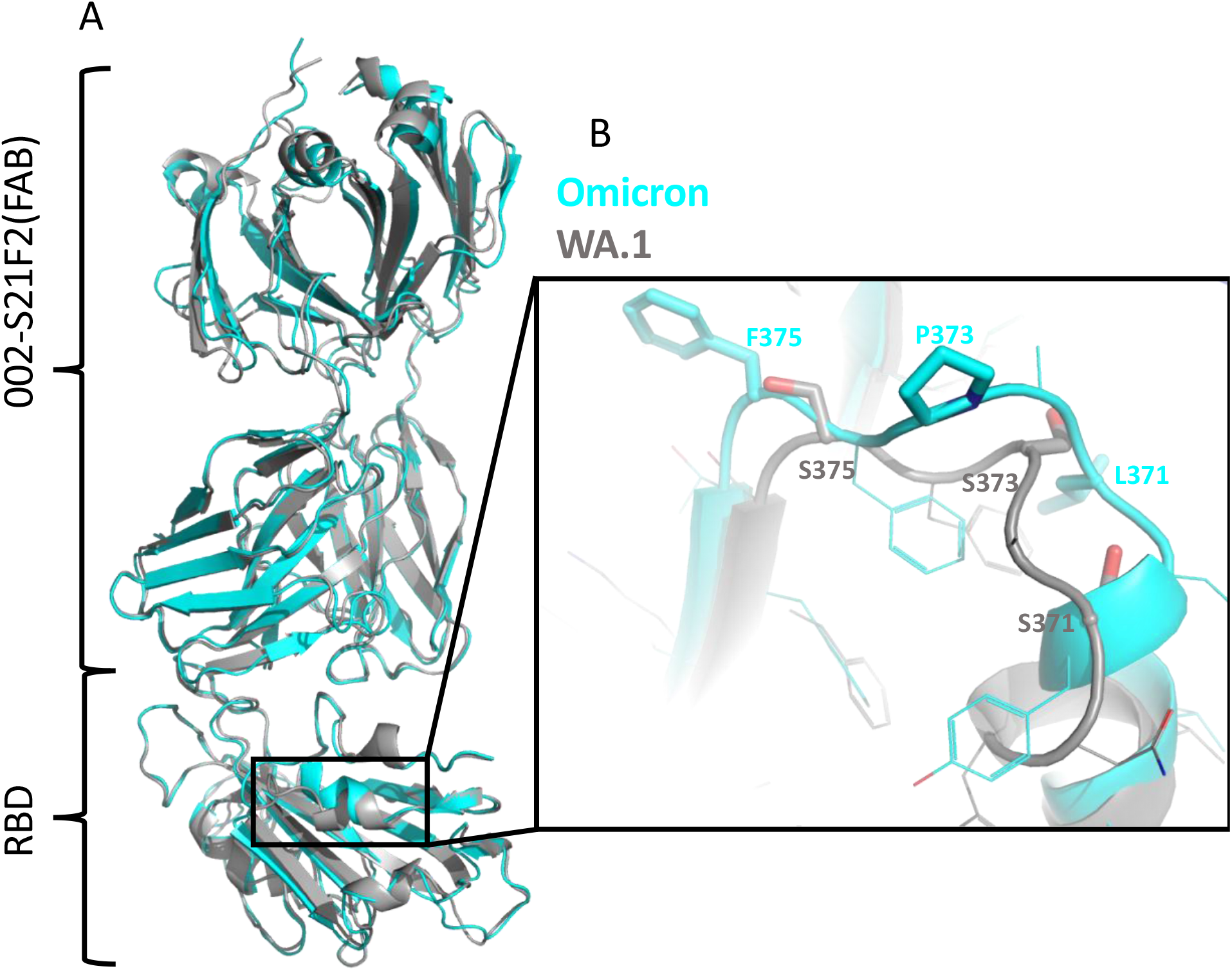
Structural comparison of WA.1 (grey) and Omicron (cyan) spike-002-S21F2 mAb complex. **(A)** Overall structural alignment of RBD and Fab region of 002-S21F2 in WA.1 and Omicron complex structures. **(B)** Zoomed in view showing the local conformation change in the specified region of RBD in WA.1 vs Omicron.

**Fig. S8.**
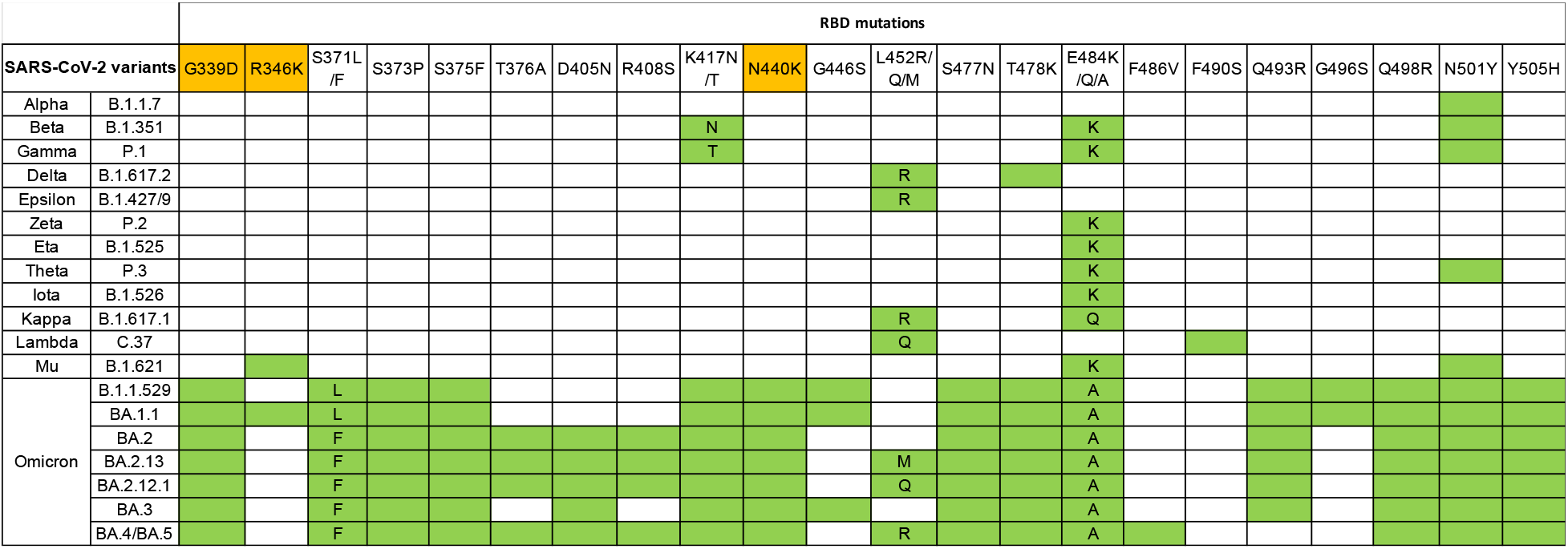
Represents the SARS-CoV-2 RBD mutations reported in key variants. The 002-S21F2 epitope residues are highlighted in orange on RBD mutations. Green color highlighted boxes show the presence of RBD mutations in each SARS-CoV-2 variant.

**Fig. S9.**
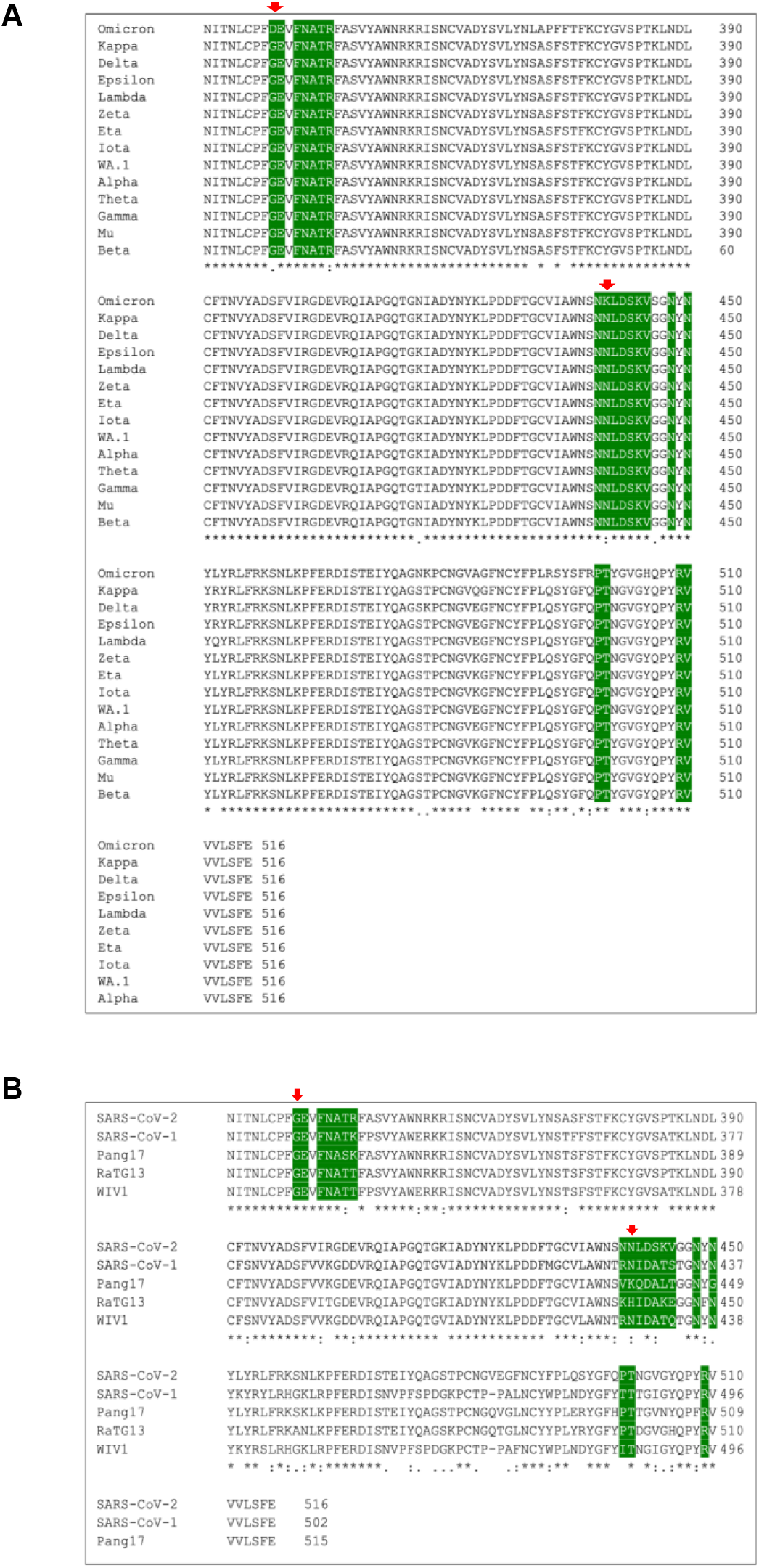
Sequence alignment of spike RBD. **(A)** from most VOC and **(B)** from different Sarbecoviruses used to derive the epitope logo. 002-S21F2 epitope residues are highlighted in green, residue. Residue positions within the 002-S21F2 epitope that are mutated in Omicron are shown as a red arrow.

**Table S1:**
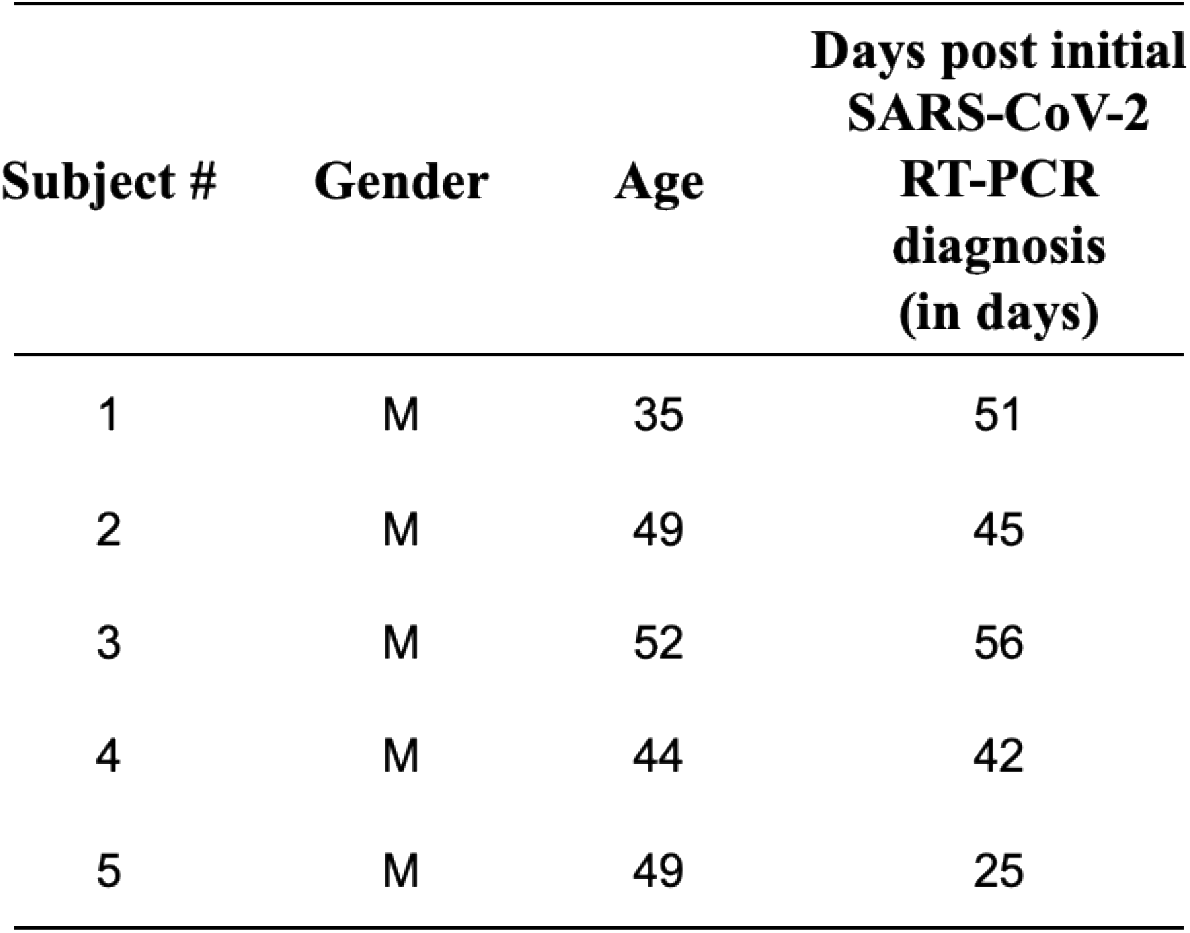
Time points chosen post initial COVID-19 diagnosis for memory B cell characterization.

**Table S2:** CryoEM data collection.

| **Spike-mAb** | **002-S21F2**<br>**WA.1** | **002-S21F2-**<br>**Omicron** |
| --- | --- | --- |
| EMDB | EMD-26262 | EMD-26669 |
| Microscope | Talos Arctica | Talos Arctica |
| Voltage (kV) | 200 | 200 |
| Detector | Gatan K3 | Gatan K3 |
| Magnification (nominal) | 79000 | 79000 |
| Energy filter slit width (eV) | 20 | 20 |
| Calibrated pixel size (Å/pix) | 1.1 | 1.1 |
| Exposure rate (e<sup>-</sup>/ Å<sup>2</sup>/sec) | 12.75 | 12.75 |
| Frames per exposure | 48 | 48 |
| Total electron exposure (e<sup>-</sup>/Å<sup>2</sup>) | 51 | 51 |
| Exposure per frame (e<sup>-</sup>/Å<sup>2</sup>) | 1.0625 | 1.0625 |
| Defocus range (µm) | 1-2.4 | 1-2.4 |
| Automation software | EPU | EPU |
| # of Micrographs used | 3863 | 4553 |
| Particles extracted | 2,249,333 | 2,553,301 |
| Particles after 2D classification | 457,277 | 310,886 |
| Total # of refined particles | 129,931 | 108,816 |
| Symmetry imposed | C1 | C1 |
| Map sharpening B-factor | 148.8 | 167.6 |
| Unmasked Resolution at 0.5/0.143 FSC (Å) | 8.0/4.0 | 8.7/4.7 |
| Masked resolution at 0.5/0.143 FSC (Å) | 4.2/3.8 | 4.5/4.1 |

**Table S3:** Model refinement and validation statistics.

| PDB | 7U0P | 7UPL |
| --- | --- | --- |
| Composition |  |  |
| Amino acids | **4102** | **4031** |
| Glycans | **32** | **54** |
| RMSD bonds (Å) | **0.004** | **0.004** |
| RMSD angles (°) | **0.605** | **0.65** |
| Mean B-factors |  |  |
| Amino acids | **102.42** | **140.31** |
| Glycans | **135.35** | **189.47** |
| Ramachandran |  |  |
| Favored (%) | **96.43** | **93.55** |
| Allowed (%) | **3.57** | **6.42** |
| Outliers (%) | **0.0** | **0.0** |
| Rotamer outliers (%) | **0.39** | **0.0** |
| Clash score | **5.24** | **7.1** |
| C-beta outliers (%) | **0.00** | **0.0** |
| CaBLAM outliers (%) | **2.93** | **3.6** |
| CC (mask) | **0.71** | **0.74** |
| CC (volume) | **0.69** | **0.73** |
| MolProbity score | **1.52** | **1.81** |

